# The influence of nutrient enrichment and fish addition on microbial communities in water and fish guts during mesocosm experiments conducted in a tropical freshwater lagoon (Lagoon Aghien, Côte d’Ivoire)

**DOI:** 10.64898/2026.10.12.758693

**Authors:** Catherine Quiblier, Marc Troussellier, Cécile Bernard, Alison Gallet, Cécile Roques, Bernard Montuelle, Éric Kouamé, Félix K. Konan, Jacques Pedron, Patrice Got, Sébastien Duperron, Jean-François Humbert, Julien C. Kalpy

## Abstract

Freshwater ecosystems in the intertropical zone have received considerably less scientific attention than those in temperate regions, with the notable exception of a few large and iconic lakes, such as Lake Victoria. Consequently, the diversity, functioning, and pollution sensitivity of microbial communities in these systems remain poorly understood. To investigate the short-term responses of microbial communities to nutrient enrichment and fish addition, we conducted *in situ* mesocosm experiments at two stations in a freshwater lagoon near Abidjan (Côte d’Ivoire). Phosphorus enrichment, either alone or in combination with nitrogen, led to a consistent increase in bacterial abundance across mesocosms, suggesting a potential phosphorus limitation of bacterial growth by these nutrients and consequently a potential competition for this resource between bacteria and microphytoplankton. In contrast, picophytoplankton abundances showed no clear or consistent response to nutrient enrichment, suggesting that other ecological processes may play a more important role in regulating these communities. High-throughput Illumina sequencing revealed contrasting patterns in the relative abundance of cyanobacterial and chloroplast-associated reads, depending on both nutrient enrichment and fish addition. These results, consistent with those obtained by cell counting in a previous study, suggest a potential top-down control of cyanobacteria by fish in Lagoon Aghien. In addition, nutrient (phosphorus and nitrogen combined) enrichment and fish addition significantly affected the amplicon sequence variant (ASV) composition of bacterial communities. Overall, these findings provide new insights into the drivers shaping microbial community structure in tropical freshwater ecosystems and are discussed in the context of previously published data on phytoplankton dynamics from the same experimental system.

## Introduction

With the goal to better understand the dynamics of cyanobacterial blooms, the relationships between bacteria (other than cyanobacteria) and phytoplankton (mostly cyanobacteria) communities have been widely studied in the past 15 years in natural freshwater ecosystems [e.g. 1-7] and also in small to medium volume experimental systems (microcosm and mesocosm) [e.g. 8-11]. All these works have permitted to describe and better understand the spatial and/or temporal dynamics of these communities under various environmental conditions and to improve our knowledge on the functional relationships that exist between members of these communities.

Most of these studies have focused on eutrophic freshwater ecosystems in temperate regions where cyanobacterial blooms frequently occur. In contrast, only a limited number of studies have been conducted in Sub-Saharan Africa, even though tropical freshwater ecosystems are known to differ markedly from their temperate counterparts due to higher water temperatures and more intense solar radiation, with consequences for their physicochemical and biological characteristics [12-15]. Moreover, all these studies consistently highlight the need for additional data on microbial communities in tropical freshwater ecosystems to improve our understanding of their functioning, particularly with respect to the relative importance of bottom-up and top-down controls, a gap that motivated the present study.

This work was conducted in the framework of an international research program (WaSAf program) dealing with the water quality of freshwater ecosystems used for the production of drinking water in three African countries (Lagoon Aghien in Ivory Coast, Lake Guiers in Senegal and Lake Victoria in Uganda). Two (Lagoon Aghien and Lake Victoria) of these three ecosystems experience recurrent cyanobacterial blooms, with negative impacts on their biological functioning and their multiple uses, including the drinking water supply [16, 17]. Within the framework of the WaSAf program, mesocosms experiments were also conducted with the aim of testing the effects of nutrient addition and fish grazing on microbial communities. In Lagoon Aghien, Ahoutou et al. [18] showed that the concomitant addition of nitrogen and phosphorus in mesocosms stimulated microphytoplankton (microalgae and cyanobacteria) growth, suggesting co-limitation by these two nutrients, whereas the addition of fish led to a marked decrease in microphytoplankton biomass and to significant changes in microphytoplankton community composition.

In the present study, we analyzed flow cytometry and high-throughput sequencing data generated from the same mesocosm experiments on bacterial and small-sized phytoplankton (picocyanobacteria and picoeukaryotes) communities, with the main objectives:

- to evaluate whether nutrient enrichment and fish addition affect the abundance and community composition of bacteria and small-sized phytoplankton, as previously found for microphytoplankton by Ahoutou et al. [18];
- to identify potential interactions, such as competition for nutrients, between all these communities;
- and finally, to gain a better understanding of the potential impact of bottom-up and top-down control on all these communities in Lagoon Aghien.

## Materials and methods

### Study site

Lagoon Aghien (5°22’N to 5°26’N and 3°49’W and 3°55’W) is a freshwater ecosystem located in the district of Abidjan in the south of the Ivory Coast. This lagoon is highly eutrophicated as shown by Ahoutou et al. [19] and used for multiple purposes (washing dishes and clothes, bathing, fishing and obtaining water for consumption) by local populations [20].

Mesocosm experiments were performed at two different stations, Débarcadère and Aghien-Télégraphe (see [18]), to detect potential spatial variations in the responses of the microbial communities to the various conditions tested. Débarcadère is a very shallow station located at the western end of the lagoon, close to the mouths of its two main tributaries (the Bété and Djibi rivers), where a water treatment plant has been constructed to supply drinking water to part of the Abidjan metropolitan area; Aghien-Télégraphe is located on the northern shore of the central part of the lagoon.

The work conducted on Lagoon Aghien was part of a research program managed in Côte d’Ivoire by the Institut Pasteur d’Abidjan, which is responsible for monitoring the water quality of the lagoon. Within this framework, no specific permit was required for the experiments. However, we met with the traditional authorities (chiefdoms) of the two villages, Aghien-Télégraphe and Débarcadère, to explain the planned experiments and obtain their oral agreement. In addition, local communities were involved in the installation of the mesocosms and were invited to visit our field laboratory. Concurrently with the mesocosm experiments, we conducted small-scale experiments using flasks in schools in both villages. These activities aimed to explain the objectives of our research to schoolchildren and to raise their awareness of pollution in the lagoon.

### Experimental setup

The mesocosm experiments were carried out in February 2018 during the long dry season. Fifteen mesocosms (200 L capacity; see the description in Ahoutou et al. [18]) were attached and filled with 160 L of lake water at Aghien-Télégraphe and Débarcadère stations. The positions of the fifteen mesocosms in the experimental setup (from M1 to M15) were randomly determined to avoid potential biases due to their position within the experimental setup (see the Figure 1 in Ahoutou et al. [18]).

**Figure 1.**
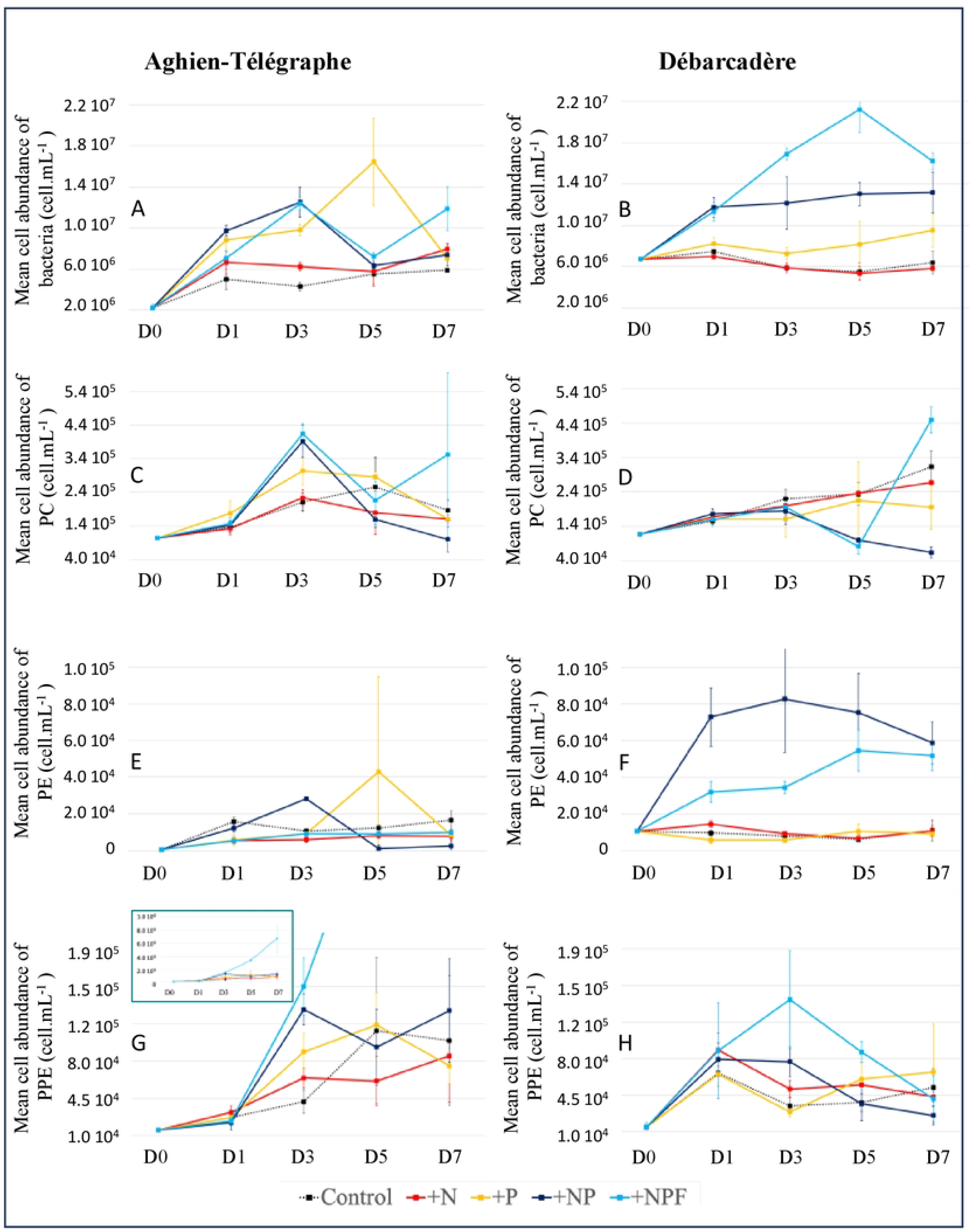
Variations in mean cell abundances (cell mL^-1^; mean values ± SD) of bacteria (A & B), phycocyanin-rich picocyanobacteria (C & D), phycoerythrin-rich picocyanobacterial (E & F) and photosynthetic picoeukaryotes (G & H) at the two stations (Aghien Télégraphe and Débarcadère). C = Control; N = +Nitrogen; P = +Phosphorus; NP = + Nitrogen and Phosphorus; NPF = + Nitrogen, Phosphorus and fish (added at day 3); D0, D1, D3, D5 & D7 = days of the experiment. PC: phycocyanin-rich picocyanobacterial; PE: phycoerythrin-rich picocyanobacterial; PPE= photosynthetic picoeukaryote

The design of the experiment at each station was as follows:

- Three control mesocosms (C1, C2 & C13) (C treatment),
- Three mesocosms (N5, N9 & N12) were supplemented only with nitrogen (16 mL of the N solution) (+N treatment), and the N-solution used for nutrient enrichment was formed from NaNO_3_ with an N concentration = 11.53 mg.mL^-1^,
- Three mesocosms (P6, P10 & P14) were supplemented only with soluble phosphorus (16 mL of the P solution) (+P treatment), and the P-solution was formed from KH_2_PO_4_ with a P concentration = 1.59 mg.mL^-1^,
- Three mesocosms (NP7, NP11 & NP15) were supplemented with nitrogen and phosphorus (16 mL of the N solution and 16 mL of the P solution) (+NP treatment),
- Three mesocosms (NPF3, NPF4 & NPF8) were supplemented with nitrogen and phosphorus (16 mL of the N solution and 16 mL of the P solution), and five Nile tilapia fish (*Oreochromis niloticus*) were added to each of them on day 3 (D3) (+NPF treatment). All fish used in the experiment were a mix of males and females; they were purchased from a fish farm located in the Lagoon Aghien. They were hand-selected with the goal of them having similar lengths and masses (∼120 mm and ∼30 g, respectively). Five control fish were measured, weighed and dissected at D3. At the end of the experiments (D7), all the fish (n= 30) were measured, weighed and dissected in accordance with ethical animal concerns and regulations. For each individual, the liver, muscles and intestines were collected, quickly frozen by using liquid nitrogen and further stored at -80 °C for cyanotoxin analysis.

For the nutrient addition, the volume of P solution added to the P, NP and NPF mesocosms was calculated by multiplying the initial soluble reactive phosphorus concentration by ten. The volume of N solution added to the N, NP and NPF mesocosms was calculated with the goal of maintaining the N/P Redfield ratio between the volume of the solutions added.

The timing of water and fish sampling in the mesocosms is presented in S1 Table.

### Water filtration and gut isolation

Raw water samples (ranging between 50–100 mL depending on the Chl-a content) were filtered on D1 for the control mesocosms and subsequently on D3 and D7 for all mesocosms. This filtration was performed through a 3.0 µm polycarbonate filter (Isopore Membrane Millipore) to concentrate the particle-associated fraction (Attached Bacteria), followed by a 0.2 µm filtration on polycarbonate filter (Nuclepore Polycarbonate Whatman) in order to concentrate the free-living fraction (Free-living Bacteria). Filters were immediately frozen at -20°C and then stored at -80°C until DNA extraction.

The fish intestines were isolated through dissection and then frozen at -20°C and then stored at -80°C.

### Cytometry analyses

Water samples (1.4 mL) were filtered through a 20 µm porosity filter to remove large particles, then fixed with prefiltered (0.2 µm) formaldehyde (2% final concentration) and stored in liquid nitrogen. All cytometry analyses were performed with a CytoFLEX flow cytometer (Beckman Coulter) Bacterial community was analyzed by using SYBER-green I (Molecular Probes) according to the protocol described by Marie et al. [21] modified with the use of a higher fluorochrome concentration by Bouvy et al. [22]. Stained bacterial cells exited at 488 nm (Blue laser 50mW) were enumerated according to their Side Scater (SSC) and the green fluorescence (FL1) using a 525/40nm filter. These cell parameters were recorded on seven decades logarithmic scale mapped onto 16 million channels. Fluorescent beads (1-2 µm, Polysciences, Warrington, PA, USA) were systematically added to each sample to estimate the relative size and nucleic acid content of bacterial cells [23]. True count (TC) beads (Becton Dickinson San Jose, CA, USA) were added to determine the volume analyzed.

Picophytoplankton (< 3µm) were analyzed after cell excitation by a red laser at 638nm (50mW), orange fluorescence (FL2, 585/42nm) for the phycoerythrin (PE), red fluorescence (FL6, 660/10nm and FL4, 690/50nm) for the phycocyanin (PC) and chlorophyll pigments respectively and then enumerated depending on the SSC and fluorescence properties (S1 Figure). Fluorescent beads (1-2 µm, Polysciences, Warrington, PA, USA) were systematically added to each sample to estimate the relative size and fluorescence intensity of picophytoplankton cells. True count (TC) beads (Becton Dickinson San Jose, CA, USA) were added to determine the volume analyzed. Data files were analyzed using the CytExpert Software (Beckman Coulter) to quantify different groups (PE, PC) of picophytoplankton cells (size and fluorescence).

### DNA extraction and PCR amplification

For water samples, DNA was extracted from the filters following Louati et al. (2015). Briefly, each filter was flash-frozen in liquid nitrogen and transferred to Lysing Matrix E tubes (MP Biomedicals, Illkirch, France) with 1.1 mL of lysis buffer (40 mM EDTA, 50 mM Tris–HCl, 0.75 M sucrose). Bead-beating was applied for 3 Å∼ 30 s at a speed of 6.5 m. s^-1^ (FastPrep1– 24, MP Biomedicals, France). Lysozyme (0.6 mg.mL^-1^) was then added to the filters and incubated at 37°C for 45 min with gentle stirring. Subsequently, sodium dodecyl sulfate (1% final concentration) and proteinase K (Thermo Scientific, France) were added, and incubated at 55C for at least 90 min. Filter-debris were pelletized by centrifugation at 14 000 g for 5 min and supernatants were collected and purified twice with phenol-chloroform-isoamyl alcohol. After precipitation with sodium acetate (0.1 vol) and cold isopropanol (0.6 vol), the nucleic acids were washed with 70% ethanol, and then re-suspended in 100 μL milliQ water. Extracted DNA was stored at -20°C until analysis.

The DNA extraction of fish gut-associated bacterial communities was performed as described by Gallet et al. [25]. Briefly, DNA was extracted from homogenized fish gut tissue (whole length, with gut content to prevent bias due to non-reproducible exclusion of gut contents) crushed using mortar and pestle to avoid the effect of longitudinal heterogeneity. The ZymoBiomics DNA mini kit (Zymo Research, Irvine, CA, USA) was used according to manufacturer’s instructions, after an initial step of cell disruption using beads (6 x 1 min).

For all samples, a 400 bp fragment of the V4–V5 region of the bacterial 16S rRNA-encoding gene was amplified using primers 515F and 926R following published PCR protocol [26]. PCR products were sequenced on an Illumina MiSeq platform (2-300 bp, paired-end sequencing, GenoToul, Toulouse, France) and paired-ends were assembled. Blank extractions and a community of known composition were used as internal controls for the sequencing procedure. The 16S rRNA gene sequence data are available in NCBI BioProject under the reference number PRJNA1451448.

### Sequence analysis

Demultiplexing of 16S rDNA gene sequences and quality control using DADA2 [27] to define ASVs were performed in QIIME2 (version 2023.5) [28]. SILVA (version 138) was used to perform taxonomic classification of each ASV [29]. The ASV feature table was rarefied to a sampling depth of 13,287 sequences per sample, the lowest number of reads found among the samples, prior to downstream analyses.

Prediction of metagenome functional content from 16S rRNA marker gene was carried out using the PICRUSt2 bioinformatics software package [30]. Briefly, 16S rRNA amplicon sequence variants (ASVs) were aligned, then gene family copy numbers of ASV inferred using reference genomes. Next, predicted gene content per ASV was calculated by using the ASV abundance table, to finally assess the metabolic pathway abundances.

### Statistical analyses

To test the impact of the five experimental treatments (Control, N, P, NP, NPF), and of the sampling time on the different measured variables, Two-way repeated measure ANOVA were performed by using PAST V5.2 Software.

Non-metric multidimensional scaling analyses (nMDS) and principal components analysis (PCA) were also performed using PAST V5.2 software. For the PCA analyses, a centered log-ratio (CLR) transformation was applied to the data prior to analysis. The nMDS analyses were based on Bray–Curtis distance matrices. Each nMDS analysis performed using PAST 5 software consisted of a sequence of 11 trials, from which the solution with the lowest stress value is selected. In the projection of the two first axes of these multivariate analyses (nMDS and PCA), convex hulls, which are the smallest convex polygons enclosing all the points belonging to each defined group, were drawn.

## Results

### Variations in the bacterial abundances during the experiments

Flow cytometry analysis showed a rapid increase in bacterial abundances in all mesocosms supplemented with at least phosphorus (+P, +NP, +NPF mesocosms) at both stations (Figure 1A & B).

At the Aghien-Télégraphe station, we found only a significant effect of Time on bacterial abundances (two-way repeated measure ANOVA; Aghien-Télégraphe: F_Time_=15.9, p=9.8×10^-9^; Débarcadère:), whereas at the Débarcadère station, significant effects on bacterial abundances were found for both Time and the Time×Treatment interaction Time (two-way repeated measure ANOVA; F_Time_=5.1, p=0.001; F_Inter._=2.7, p=0.0005).

All bacterial cytometry data are available in S2 Table.

### Variations in the picocyanobacterial and photosynthetic picoeukaryotes abundances during the experiments

The cytometry analysis of the picocyanobacterial communities allowed us to distinguish phycocyanin-rich picocyanobacteria (PC) from phycoerythrin-rich picocyanobacteria (PE), the former being the most abundant (S1 Figure). As shown in Figure 1C-D, no clear trend was observed in the variation of PC cell abundances over the course of the experiment at either station but it is worth noting that the temporal evolution of PC cell abundances in the NP and NPF mesocosms was highly similar at each station. In particular, the NPF mesocosms exhibited a sharp increase in PC cell abundances at D7, whereas no such increase was observed in the NP mesocosms on the same day. At the Aghien-Télégraphe station, only a significant effect of Time was found (two-way repeated measure ANOVA; F_Time_=15.9, p=9.7×10^-9^), whereas at the Débarcadère station, significant effects in the PC cell abundances were found for Time (two-way repeated measure ANOVA; F_Time_=7.6, p=5.6×10^-5^) and for the Time×Treatment interaction (two-way repeated measure ANOVA; F_Inter._=2.4, p=0.003).

Phycoerythrin-rich picocyanobacteria (PE) were much less abundant than PC picocyanobacteria at the beginning of the experiment (10^2^-10^4^ cells.mL^-1^ *versus* 10^5^ cells.mL^-1^) (Figure 1E&F). While there was no clear trend in the temporal variation of PE cell abundances at the Aghien-Télégraphe station, a strong increase in PE cell abundances was observed in the NP and NPF mesocosms at the Débarcadère station (Figure 1E & F). No significant effects of Treatment, Time or the Time×Treatment interaction were found for PE cell abundances at Aghien-Télégraphe (two-way repeated-measures ANOVA), whereas significant effects of Time and the Time×Treatment interaction were found at Débarcadère (two-way repeated measure ANOVA with F_Time_=4.0, p=0.007; F_Inter._=2.3, p=0.003).

Finally, no consistent trend was observed in the variation of photosynthetic picoeukaryote (PPE) abundances across the different mesocosms at either station (Figure 1G-H). The most pronounced temporal shift was a sharp increase in cell abundance within the NPF mesocosms at the Aghien-Télégraphe station. At the Aghien-Télégraphe station, significant effects on PPE abundances were found for both Time and the Time×Treatment (two-way repeated-measures ANOVA; F_Time_=7.0, p=0.0001; F_Inter_.=2.3, p=0.003), whereas at the Débarcadère station, only a significant effect of Time was found (two-way repeated ANOVA; F_Time_=12.4, p=2.8×10^-7^).

All cytometry data on picocyanobacteria and picophytoplankton are available in S3 Table.

### Variations in the in the number of 16S rRNA gene reads assigned to cyanobacteria and chloroplasts during the experiment

As expected, more cyanobacterial and chloroplast reads were recovered in the 3.0 µm fraction than in the 0.2 µm fraction at both stations (Figure 2). Additionally, highly contrasting responses were observed between cyanobacteria and chloroplasts under the NP and NPF treatments at D3 and D7. Cyanobacterial read counts were highest in the NP3, NP7, and NPF3 mesocosms and lowest in the NPF7 mesocosms. In contrast, chloroplast reads showed the opposite pattern, with low counts in the NP3, NP7, and NPF3 mesocosms and the highest counts in the NPF7 mesocosms (see the two red dashed rectangles in Figure 2).

**Figure 2.**
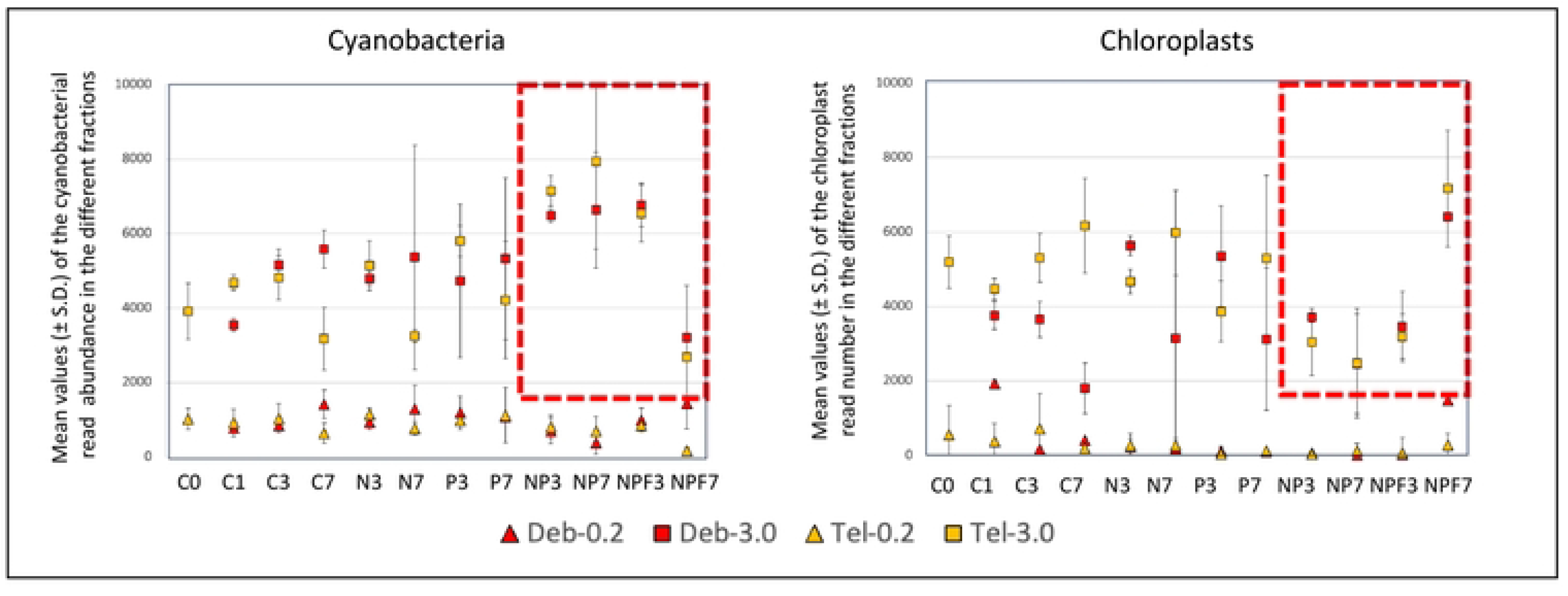
Variations in the mean number (± standard deviation, SD) of cyanobacterial and chloroplast 16S rRNA reads in the two water filtered fractions (0.2 and 3.0 µm) over the course of the experiments (three mesocosms per treatment) performed at the two experimental stations (Tel: Aghien-Télégraphe; Deb: Débarcadère). C = Control; N = +Nitrogen; P = +Phosphorus; NP = + Nitrogen and Phosphorus; NPF = + Nitrogen, Phosphorus and fish (added at day 3); 0, 1, 3 & 7 = days of the experiment. No water samples were collected from the water used to fill the mesocosms on day 0 at the Débarcadère station, whereas three water samples were collected at the Aghien-Télégraphe station (C_0_ value in the figure).

### Variations in the richness and diversity of the bacterial communities in water and in fish gut during the experiment

All 16S rRNA sequencing data are available in S4 Table. Bacterial reads (excluding cyanobacteria) were the most abundant in the 16S rRNA-encoding gene Illumina dataset (53.6%), followed by chloroplast reads (26.0%), cyanobacterial reads (19.8%) and other reads (Archaea and unclassified reads, 0.6%).

The average ASV richness was significantly higher in water (260±45) than in fish gut (180±39) (Student t-test, p<0.0001), while there was no significant difference in the ASV diversity of these communities as estimated by the Shannon index (4.6±0.2 in water and 4.7±0.3 in gut fish).

In the water microbial communities, there was a significant difference (paired t-test, p=0.001) in the ASV richness (and in Chao1 index) between the 0.2 µm fraction (mean ASV richness = 287.4±44.5) and the 3.0 µm fraction (mean ASV richness=249.2±40.6) at the Aghien-Télégraphe station but not at the Débarcadère station (mean ASV richness=249.9±41.0 for the 0.2 fraction and 244.3±33.8 for the 3.0 fraction). For ASV diversity (Shannon index), a significant difference (paired t-test, p<0.0001) was also found at the Aghien-Télégraphe station between the 0.2 µm fraction (average diversity=4.71±0.22) and the 3.0 µm fraction (mean ASV diversity = 4.51±0.17) but not at the Débarcadère station (mean ASV diversity = 4.67±0.22 for the 0.2 µm fraction and 4.57±0.14 for the 3.0 µm fraction).

As shown in Supplemental Figure 2, only moderate changes occurred over the course of the experiment in the richness and diversity of the microbial communities. No significant effects of Treatment, Time or the Time×Treatment interaction (two-way repeated-measures ANOVA) were found for ASV richness or diversity (Shannon-index) in either water fractions (0.2 µm and 3.0 µm) at Aghien-Télégraphe, and in the 3.0 µm fraction at Débarcadère. At this latter station, in the 0.2 µm fraction, a significant effect of the Time×Treatment interaction was found for ASV richness (two-way repeated measure ANOVA with F_Inter._=2.7, p=0.04) and a significant effect of Time was found for ASV diversity (two-way repeated measure ANOVA with F_Time_=7.0, p=0.02).

For the microbial communities collected from fish guts, no significant differences in the ASV richness and diversity were found between control fish, Aghien-Télégraphe fish and Débarcadère fish (Mann-Withney U-test).

### Variations in the structure and composition of the microbial communities during the course of the experiments

#### Cyanobacterial communities in water and fish

Figure 3 shows the variations in the proportions of 16S rRNA gene reads assigned to different cyanobacterial families during the experiment in the 0.2 and 3.0 µm fractions. Very similar patterns were observed in the 3.0 µm fraction at the two study stations (Aghien-Télégraphe and Débarcadère). In the C and N mesocosms, most reads were assigned to the *Microcystaceae* and *Phormidiaceae* families throughout the experiment. With the exception of the NPF mesocosms at D7, the *Nostocaceae* family, along with *Microcystaceae* and *Phormidiaceae* to varying extents, accounted for the highest number of reads in the P, NP, and NPF mesocosms. At D7 in the NPF mesocosms, most reads were assigned to the *Microcystaceae* family, predominantly to the genus *Microcystis*, concomitant with a sharp decrease in cyanobacterial read abundance.

**Figure 3.**
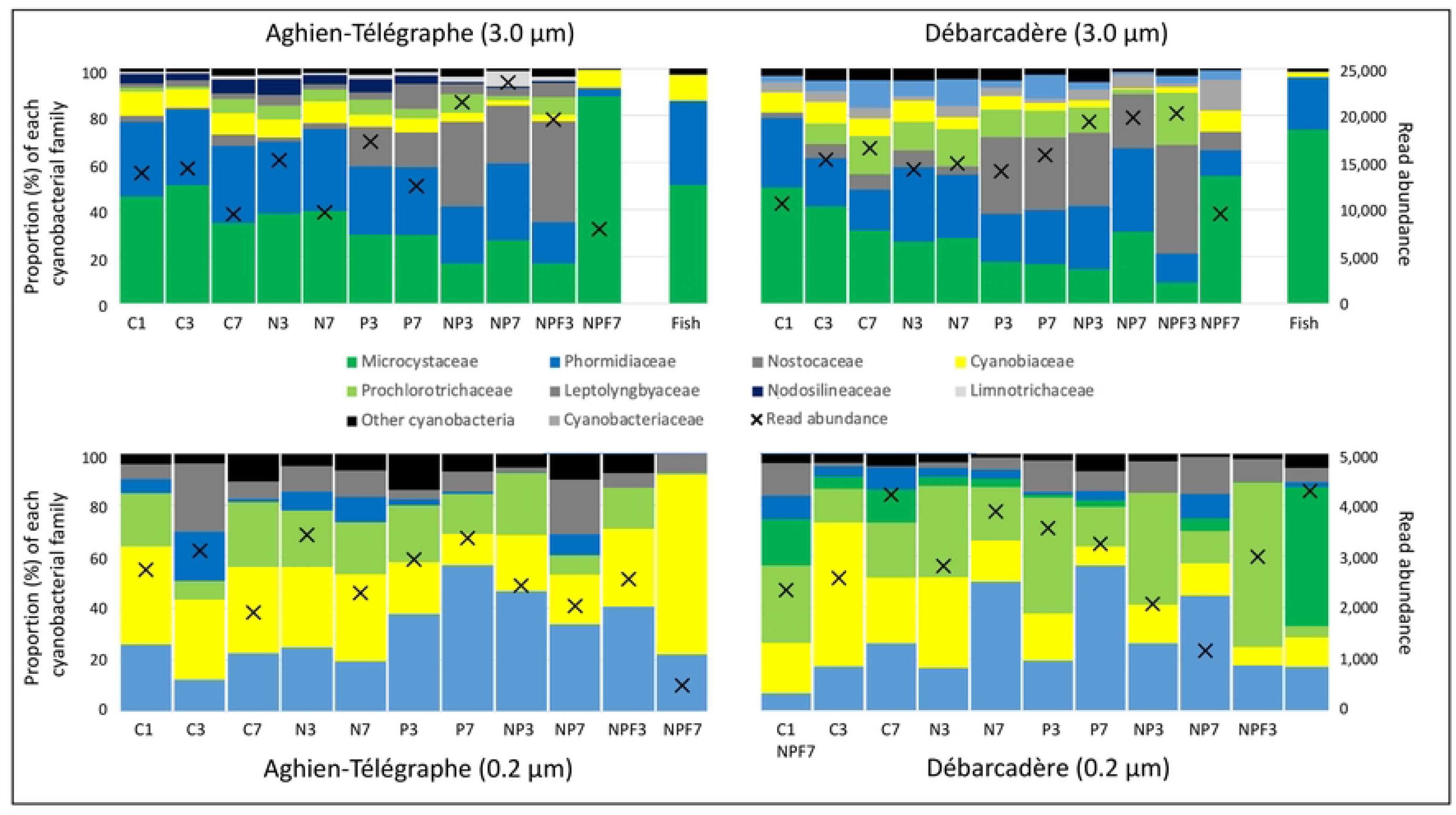
Variations in total cyanobacterial 16S rRNA gene read abundance and in the relative abundance of the main cyanobacterial families in the two filtered water fractions (0.2 and 3.0 µm) and in fish guts throughout the experiments at both stations (Aghien-Télégraphe and Débarcadère). C = Control; N = +Nitrogen; P = +Phosphorus; NP = + Nitrogen and Phosphorus; NPF = + Nitrogen, Phosphorus and Fish (added at day 3 after the sampling in the mesocosms); Fish = fish gut; 1, 3, 7 = days of the experiment

In the 0.2 fraction, most of the reads were assigned to *Leptolynbyaceae*, *Cyanobiaceae* and *Prochlorotrichaceae* families, with variations in their relative proportions depending on the station (Aghien-Télégraphe and Débarcadère) and the treatment. As in the 3.0 µm fraction, the composition of the cyanobacterial communities in NPF mesocosms at D7 was very different from that observed in the other treatments and dates.

Finally, the cyanobacterial communities found in the fish guts at the two stations (Aghien-Télégraphe and Débarcadère) were similar to those found in the 3.0 µm fraction of the C and N mesocosms, with most reads assigned to the *Microcystaceae* family (mainly *Microcystis* genus) and, to a lesser extent, to the *Phormidiaceae* family.

#### Microalgae communities in water and fish

The microalgal communities in the mesocosm water were characterized using chloroplast-assigned 16 rRNA sequences. This analysis was performed only in the 3.0 fraction at both stations (Aghien-Télégraphe and Débarcadère) due to the very low number of chloroplast reads in the 0.2 fraction. As shown in S3 Figure, approximately 50% of the reads were assigned to the *Xanthophyceae* class and around 30% corresponded to unidentified chloroplast sequences. Finally, the proportion of reads assigned to the *Chlorophyceae* class varied from a few percent to 20%. The most notable differences among the mesocosms were observed in the NPF treatment on D7, which showed the lowest proportion of reads from the *Xanthophyceae* class and the highest proportion from the *Chlorophyceae* class. Lastly, the microalgal communities found in the gut of fish in the NPF mesocosms at both stations closely resembled those in the NPF mesocosm water on D7, except for a higher proportion of *Chlorophyceae* reads.

#### Bacterial communities (excluding cyanobacteria) in water and fish

At the phylum level, in the NPF mesocosms at D7 and across both stations, most water bacterial reads (excluding cyanobacteria) were assigned to the Pseudomonadata (Proteobacteria) (34%), Bacteroidota (33%), Actinomycetota (17%), and Verrucomicrobiota (8%) phyla (Supplemental Figure 4). In contrast, in fish guts, most bacterial reads at D7 and across both stations were assigned to the Bacillota (Firmicutes) (37%), Planctomycetota (17%), Fusobacteriota (Fusobacteria) (13%), and Verrucomicrobiota (11%) phyla (S4 Figure). Similarly, at the ASV level, non-metric multidimensional scaling analyses revealed a clear distinction between the bacterial communities of fish guts and those of water samples at both stations (S5 Figure).

Four nMDS analyses were performed on the bacterial communities of each water filtered fraction (0.2 µm and 3.0 µm) at the two experimental stations (Figure 4). Stress values of the nMDS analyses were notably high across all analyses (ranging from 0.20 to 0.55), indicating that the resulting ordinations provided a poor representation of the data in reduced dimensional space. For this reason, only the 0.2 µm fraction was retained for further analysis at both stations, as it yielded the lowest stress values (0.2 for Aghien-Télégraphe and 0.3 for Débarcadère). A consistent pattern was observed along the first axis of the analysis: C mesocosms clustered on the negative side, NP and NPF mesocosms were located on the positive side, with the N and P mesocosms positioned intermediately between them.

**Figure 4.**
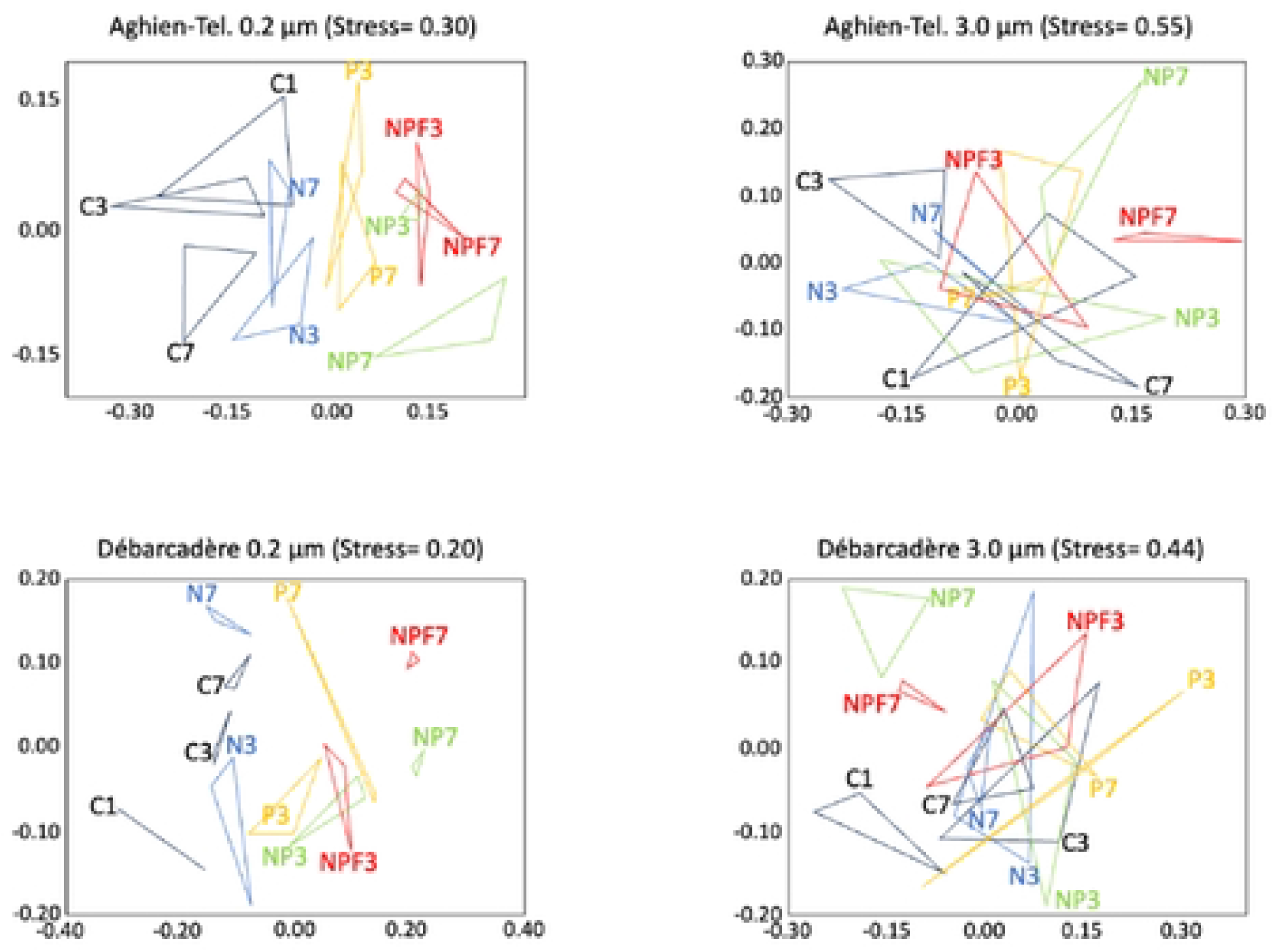
Non-metric multidimensional scaling (nMDS) biplots based on Bray–Curtis dissimilarity matrices of ASV abundances in bacterial communities (excluding cyanobacteria) in the two filtered water fractions (0.2 and 3.0 µm) at the two experimental stations (Aghien Télégraphe and Débarcadère). C = Control; N = +nitrogen; P = +phosphorus; NP = + nitrogen and phosphorus; NPF = + nitrogen, phosphorus and fish (added at day 3 after the sampling in the mesocosms); Fish = fish gut; 1, 3, 7 = days of the experiment,

### PICRUSt analysis performed of the bacterial communities in the two water filtered fractions (0.2 and 3.0 µm)

Very similar results were obtained at the two stations (Aghien-Télégraphe and Débarcadère) in the principal component analyses (PCA) of the PICRUSt-predicted functional profiles of bacterial communities in the two filtered water fractions (0.2 and 3.0 µm) (Figure 5) (S5 Table). First, there was a clear distinction between the functional profiles of these two filtered fractions in the PCA projection onto the first two components. Second, for both fractions, the functional profiles of the Control (C) mesocosms were clearly separated from those of the NP and NPF mesocosms, while those of the N and P mesocosms occupied an intermediate positions.

**Figure 5.**
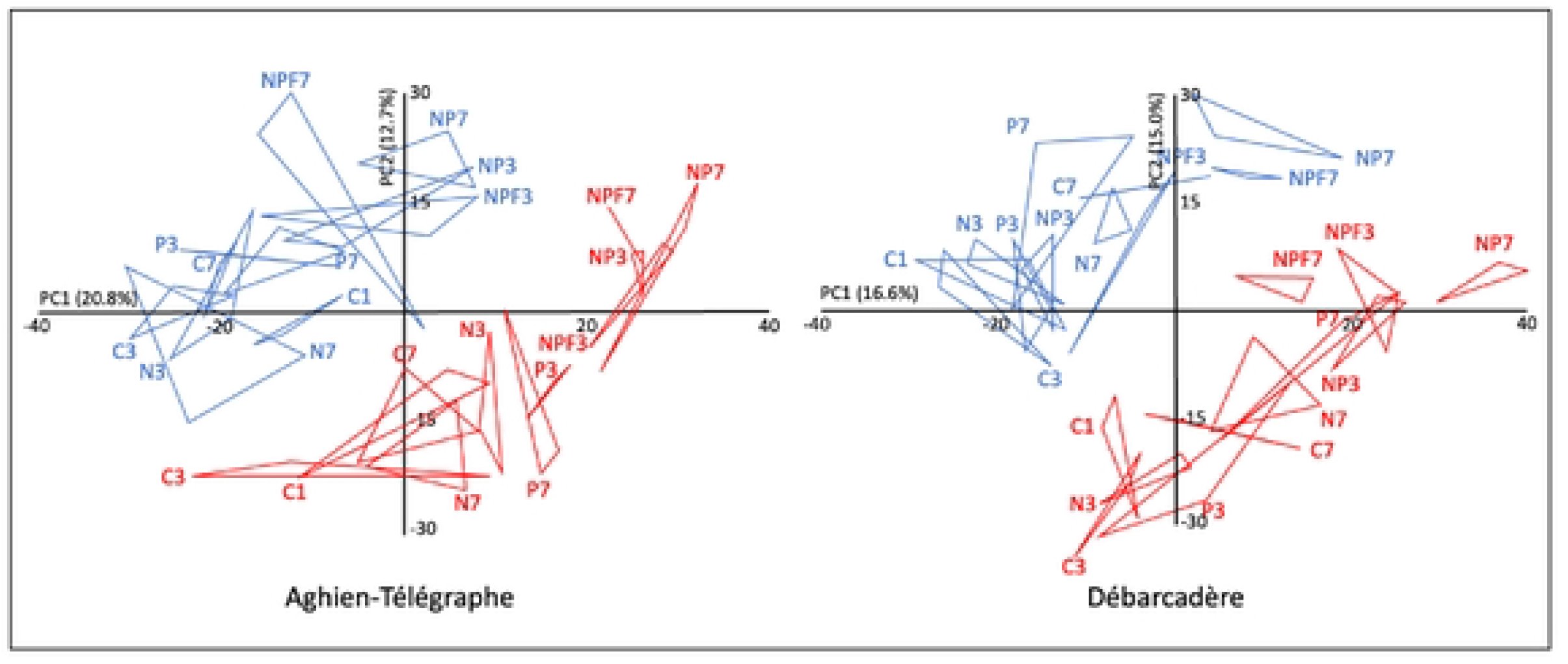
Principal component analyses performed on PICRUSt data from the 0.2 and 3.0 µm filtered water fractions over the course of the experiment (D1, D3, and D7) at Aghien– Télégraphe (A) and Débarcadère (B) stations. The convex hulls (red for the 0.2 µm fraction and blue for the 3.0 µm station) encompass the points obtained for each treatment (C, N, P, NP, NPF) and sampling day (1, 2, and 3). C = Control; N = +Nitrogen; P = +Phosphorus; NP = +Nitrogen and Phosphorus; NPF = + Nitrogen, Phosphorus and Fish (added at day 3 after the sampling in the mesocosms).

Examination of the correlations between variables (metabolic pathways) and PC scores in the PCA, indicated that, among the pathways strongly positively or negatively correlated with PC1 & 2, several are involved in the biosynthesis of isoprenoid quinones (e.g. PWY-5838, 5845, 5862, 5861, 5862, 5897; 5898, 5898 for the PC1 and PWY-5855, 5856, 5857 and 6708 for the PC2).

## Discussion

In the paper by Ahoutou et al. [18], dealing with the same experiment and reporting data obtained on cyanobacteria and microalgae by cell counting under the microscope, it was shown that the addition of both nitrogen and phosphorus stimulated phytoplankton growth and resulted in significant shifts in the diversity and composition of these communities. Furthermore, the introduction of fish led to a marked decrease in phytoplankton biomass and induced a shift in community structure from cyanobacterial to chlorophyte dominance, particularly at the Aghien-Télégraphe station [18]. Direct comparison between the phytoplankton communities data presented in Ahoutou et al. [18] and those of the present study is challenging, as the former were expressed in biovolume units, whereas the latter are reported as numbers of Illumina sequencing reads. Moreover, in the present study, microalgal community composition was assessed using 16S plastid sequences, which are not considered the best markers for microalgal communities (e.g. Kezlya et al. [31]). However, these and other ones [32, 33] have shown that environmental 16S rRNA datasets can yield useful information on eukaryotes, albeit with lower sensitivity and taxonomic resolution than eukaryotic markers. Nevertheless, both approaches (biovolume estimation and 16S rRNA gene marker analysis) converged in highlighting a significant impact of fish addition in the NPF mesocosms at both Aghien-Télégraphe and Débarcadère stations. This effect resulted in a decrease in cyanobacterial read numbers in all NPF7 mesocosms, accompanied by an increase in chloroplast read numbers. As previously emphasized in Ahoutou’s study, this finding aligns with the work of Torres et al. [34], which demonstrated that omnivorous filter-feeding fish like the Nile tilapia (Oreochromis niloticus) have the potential to control cyanobacterial populations in eutrophic tropical lakes. As an alternative to the hypothesis of a differential impact of fish grazing on cyanobacteria and chlorophytes to explain the contrasting variations in their read numbers in the NPF mesocosms, it has been shown also that fish excretion can supply nutrients (N and P) to primary producers, and that some phytoplankton taxa respond more to nutrient excretion than to direct herbivory, while grazing is more important for others (see the review by Vanni [35]). Finally, Zeng et al. [36] found that Nile tilapia feeding and defecation favor a community shift from Cyanobacteria to Chlorophyta, these latest being more resistant to the passage through tilapia gut, and a similar finding was also reported by McDonald [37] for the blue tilapia (*Oreochromis aureus*).

Expanding upon the findings of Ahoutou et al. [18], the present study provides data on both pico-phytoplankton and bacterial communities. Among the picocyanobacteria, phycocyanin-rich (PC) cells were more abundant than phycoerythrin-rich (PE) cells at the start of the experiment, just after the mesocosm were filled with lake water. This result was anticipated, given the eutrophic status of Lagoon Aghien [19] and previous findings indicating that eutrophic conditions, in particular light limitation, tend to favor PC-dominant picocyanobacterial populations [38].

Temporal variations in PC, PE, and total picophytoplankton cell abundances during the mesocosm experiments differed markedly between the two stations, as illustrated, for example, by the PE community, for which there was a clear impact of N and P addition on cell abundance at the Débarcadère station but not at Aghien-Télégraphe. In the same way, very contrasted responses were observed at D5 and D7 after the addition of fish in the mesocoms. These findings differ markedly from those obtained in the same experiment by Ahoutou et al. [18] on large phytoplankton communities for which there was a clear impact of N and P addition and of fish addition on these communities at the Aghien-Télégraphe and Débarcadère stations.

Taking all these elements into account, the very high variability observed in the responses of the different picophytoplanktonic communities to nutrient and fish additions among the different mesocosm conditions, as well as between the two experimental stations, strongly suggests that additional processes, which were not investigated in the present study, may also play an important role in determining the dynamics of these communities. Among these potentially important processes, zooplankton grazing would have been particularly relevant to investigate, as it may exert a strong influence on picophytoplankton communities through top-down control [e.g., 39, 40]. Unfortunately, we were unable to assess the contribution of zooplankton grazing in the present study.

Regarding heterotrophic bacterial communities (excluding cyanobacteria), our flow cytometry data showed a significant increase in bacterial cell abundances in the NP and NPF mesocosms at both stations, as well as in the P mesocosm at Aghien-Télégraphe and, to a lesser extent, at Débarcadère. The existence of a phosphorus limitation of bacterial growth, coupled or not to a nitrogen and/or an organic carbon limitation, has already been described in several temperate and cold lakes and ponds [e.g. 42-45] but more rarely in tropical ones [46, 47]. Knowing that Ahoutou et al. [18] also revealed phosphorus and nitrogen limitation of phytoplankton, this suggests that potential competition, at least for phosphorus, might occur under limiting conditions between bacteria and microbial primary producers in Lagoon Aghien, as has already been observed in other freshwater ecosystems [e.g. 48, 49].

In addition to changes occurring in their abundances, we found also changes in the structure and composition of microbial communities during the experiments. In cyanobacterial communities, the addition of phosphorus resulted mainly in a decrease of the number of reads belonging to *Microcystaceae* (mainly *Microcystis* genus) coupled to an increase of reads belonging to *Nostocaceae* (mainly *Raphidiopsis* and *Dolichospermum* genera) in P, NP and NPF mesocosms at the two stations. These results are consistent with the findings of Rigosi et al. [50] and Wan et al. [51], who reported that diazotrophic genera such as *Dolichospermum* are more effective competitors than the *Microcystis* genus under high phosphorus availability, whereas the opposite is true under low phosphorus availability. Interestingly, we found also that the addition of fish on D3 in the NPF mesocosms resulted at day 7, in a dominance of *Microcystaceae* reads within the cyanobacterial communities and to a strong decrease in the total number of cyanobacterial reads compared to NP3, NPF3 and NP7 mesocosms. This suggests that inside the cyanobacterial communities, filamentous genera from the *Phormidiaceae* and *Nostocaceae* families were either preferentially grazed by fish or *Microcystaceae* cells were more resistant to digestion than *Phormidiaceae* and *Nostocaceae* cells in the fish’s intestines. However, this latter hypothesis is not supported by the literature. Jancula et al. [52] and Wang et al. [53] reported that *Microcystis* cells were not viable after passing through the digestive tract of *O. niloticus*, while the opposite result was observed for silver carp.

As expected, a clear distinction was observed between bacterial communities (excluding cyanobacteria) associated with fish guts and those present in the water column. This finding is consistent with the results of Gallet et al. [25], who compared waterborne and fish gut-associated bacterial communities in Lagoon Aghien, as well as with many other studies showing that fish gut microbiota are typically dominated by members of the Bacillota, Pseudomonadota, and Fusobacteriota phyla [e.g. 54-56]. Notably, at the Aghien-Télégraphe station, the waterborne bacterial community in the 3.0 µm fraction from the NPF mesocosms on D7 clustered closely with the fish gut-associated communities in our nMDS analysis. This finding suggests that fish have dramatically impacted the attached bacterial community in these mesocosms, and further supports the hypothesis that fish may act as significant reservoirs of bacterial diversity in aquatic ecosystems, as previously proposed by Troussellier et al. [57] and Gallet et al. [25].

Among the water-associated communities, additional distinctions were evident between the 0.2 µm and 3.0 µm size fractions. Firstly, the convex hull areas were larger in the 3.0 µm fraction compared to the 0.2 µm fraction, indicating greater heterogeneity in the composition of bacterial communities among replicates in the 3.0 µm fraction. Moreover, no clear effect of the treatment was detected in the composition of these communities (overlay of their convex hulls), except for NP7 and NPF7 mesocosms at both stations. These findings are particularly relevant in light of the results reported by Ahoutou et al. [18] on the temporal evolution of phytoplankton community composition in the same mesocosms. Indeed, their study revealed considerable heterogeneity in community structure, as evidenced by substantial overlap among the convex hulls in the PCA analysis. Moreover, the phytoplankton communities in the NP7, and even more so in the NPF7 mesocosms, were the most distinct from the others, highlighting the strong influence of nutrient and fish additions on these communities. Taken together, these results strongly support the close relationships between particle-attached bacterial communities and phytoplankton communities, as previously observed in numerous field studies conducted in both temperate and tropical environments [e.g. 58, 59]. Finally, our results show that the addition of phosphorus, and even more so the combined addition of phosphorus and nitrogen, significantly influenced the composition of bacterial communities in the 0.2 µm fraction. This observation is consistent with cytometry-based analyses, which revealed stimulated bacterial growth following nutrient enrichment, particularly with phosphorus alone and in combination with nitrogen. The 0.2 µm fraction is generally considered to represent the free-living bacterial community. Our findings suggest that these free-living bacteria should not be viewed solely as a passive reservoir for the recruitment of attached bacteria, as previously suggested by Louati et al. [6].

Finally, the PICRUSt analysis on bacterial communities at both stations revealed a clear separation between the 0.2 µm and 3.0 µm fractions and, within both fractions, between the C mesocosms and the NP and NPF mesocosms, with the N and P mesocosm positioned intermediately. When looking at the metabolic pathways strongly associated with the PC1 and PC2 axes, several of them concerned the biosynthesis of isoprenoid quinones, including menaquinones and ubiquinones. Isoprenoid quinones play a central role in bioenergetics, serving as essential electron and proton carriers in the respiratory chains of most organisms. Menaquinones (MK) are the most widespread respiratory quinones among microorganisms, being present in many groups of Archaea and Bacteria. In contrast, ubiquinones (UB) are restricted to α-, β-, and γ-Pseudomonadota, as well as eukaryotes [60]. Notably, almost all cyanobacteria lack both UB and MK [60, 61]. Furthermore, it has been shown that the biosynthetic pathways of MK and UB are evolutionarily related [62]. Taken together, these findings suggest that the PICRUSt-based analyses support our observation that the different conditions tested in the mesocosms affect the composition of the bacterial communities, rather than revealing functional differences in their metabolic capabilities.

In conclusion, these findings corroborate and complement those reported by Ahoutou et al. [18] and further highlight the relevance of mesocosm approaches for advancing our understanding of the biological functioning of tropical lake ecosystems. In particular, mesocosm experiments provide a valuable framework for investigating the effects of key ecological processes, such as nutrient enrichment and fish grazing, as well as their potential interactive effects on ecosystem biodiversity and functioning. By extending the findings previously reported by Ahoutou et al. [18], the present study further underscores the importance of considering a broad range of microbial community components to gain a comprehensive understanding of the direct and indirect effects of eutrophication and fish addition on microbial communities. One limitation of the present study is the lack of data on zooplankton, although its characterization was initially planned, it could ultimately not be carried out. This represents an important gap in our assessment, as several findings suggest that zooplankton grazing and associated trophic cascading effects in the fish-amended mesocosms may have contributed to some of the differences observed between the two experimental stations. Incorporating zooplankton dynamics into future experiments would therefore provide a more complete understanding of the mechanisms underlying the observed responses of microbial communities to nutrient enrichment and fish addition. Conversely, a major strength of this study lies in the diversity and complementarity of the approaches employed, including microscopic counting, flow cytometry, and high-throughput sequencing. By integrating and cross-validating results obtained using these complementary methods, our approach provides a more comprehensive characterization of microbial communities while helping to overcome some of the limitations inherent to each individual technique. This multi-method approach therefore provides a stronger basis for assessing the complex and potentially indirect effects of environmental perturbations on microbial community structure and dynamics in tropical lake ecosystems.

## Acknowledgements

The authors are extremely grateful to the inhabitants and chiefdoms of the two villages (Aghien-Télégraphe and Débarcadère) for their agreement to work on the lagoon and their help during the experiment.

**S1Table.** Sampling schedule in the mesocosms for flow cytometry analyses and 16S rRNA gene sequencing (water and fish).

**S2 Table.** Bacterial cytometry data collected at the two stations, Aghien Télégraphe and Débarcadère.

**S3 Table.** Cytometry data on picocyanobacteria and picophytoplankton samples collected at the two stations, Aghien Télégraphe and Débarcadère.

**S4 Table.** 16S rRNA sequencing data from water samples and fish samples collected at the two stations, Aghien-Télégraphe and Débarcadère.

**S5 Table.** Data obtained by the PICRUSt analysis performed on bacterial 16S rRNA sequences at the two stations, Aghien-Télégraphe and Débarcadère.

**S1 Figure.** Examples of cytograms (dot plots) of phototrophic planktonic populations. **A.** Total picoplankton (pico) and nanoplankton (>3 µm, Nano H & Nano B) populations after blue laser excitation (488 nm), collected on red fluorescence FL4H (690/50 nm) and Side Scatter (SSC). **B.** Phycoerythrin-containing (PE) cell populations after blue laser excitation (488 nm), collected on orange fluorescence FL2H (585/42 nm) and Side Scatter (SSC). **C.** Phycocyanin-containing (PC1-PC2-PC3-PC4) cell populations after red laser excitation (638 nm), collected on orange fluorescence FL6H (660/10 nm) and Side Scatter (SSC). **D.** 1-2 µm fluorescent bead standards and TruCount (TC) beads for fluorescence calibration (FL1 at 525 nm), relative size (SSC), and volumetric measurement (TC), after blue laser excitation (488 nm).

**S2 Figure.** Variations in the richness and diversity in the two water filtered fractions (0.2 and 3.0 µm) during the course of the experiments performed at the two experimental stations (Aghien-Télégraphe and Débarcadère). C = Control; N = + Nitrogen; P = + Phosphorus; NP = + Nitrogen and Phosphorus; NPF = + Nitrogen, Phosphorus and Fish (added at day 3 after the sampling in the mesocosms); 0, 1, 3, 7 = days of the experiment.

**S3 Figure.** Variations in the proportions of the different classes of microalgae found in the 3.0 µm water filtered fraction by the analysis of 16S rRNA chloroplast sequences during the mesocosm experiments performed at the two experimental stations (Aghien-Télégraphe and Débarcadère). C = Control; N = +Nitrogen; P = +Phosphorus; NP = + Nitrogen and Phosphorus; NPF = + Nitrogen, Phosphorus and Fish (added at day 3 after the sampling in the mesocosms); Fish = fish gut; 1, 3, 7 = days of the experiment

**S4 Figure.** Phylum-level composition of bacterial communities (excluding cyanobacteria) in water and fish gut sampled from the NPF mesocosms on day 7 (both stations combined).

**S5 Figure.** Non-metric multidimensional scaling (nMDS) biplots based on Bray–Curtis dissimilarity matrices of the ASV abundances in the bacterial communities (excluding cyanobacteria) in the two water filtered fractions (0.2 and 3.0 µm) and fish gut at the two experimental stations (Aghien Télégraphe and Débarcadère). C = Control; N = +Nitrogen; P = +Phosphorus; NP = + Nitrogen and Phosphorus; NPF = + Nitrogen, Phosphorus and Fish (added at day 3 after the sampling in the mesocosms); 1, 3, 7 = days of the experiment; Fi-Cont = bacterial communities from the guts of control fishes; Fi-MX = bacterial communities from the guts of fishes in the M3, M4 and M8 mesocosms. Red convex hulls = Bacteria in 3.0 µm fraction; Blue convex hulls = Bacteria in 0.2 µm fraction; Dashed lines = control mesocosm; bold line and character = NP and NPF mesocosms. One fish was removed from the analysis at Aghien-Télégraphe station in mesocosm 8 due to incoherent data.

**S1 Information.** Inclusivity questionnaire.

